# Activity and function of the endothelial sodium channel is regulated by the effector domain of MARCKS like protein 1 in mouse aortic endothelial cells

**DOI:** 10.1101/2024.06.25.600595

**Authors:** Ling Yu, Niharika Bala, Van-Anh L. Nguyen, Leah Kessler, John F. LaDisa, Abdel A. Alli

## Abstract

The endothelial sodium channel (EnNaC) plays an important role in regulating vessel stiffness. Here, we investigated the regulation of EnNaC in mouse aortic endothelial cells (mAoEC) by the actin cytoskeleton and lipid raft association protein myristoylated alanine-rich C-kinase substrate like protein 1 (MLP1). We hypothesized that mutation of specific amino acid residues within the effector domain of MLP1 or loss of association between MLP1 and the anionic phospholipid phosphate PIP2 would significantly alter membrane association and EnNaC activity in mAoEC. mAoEC transiently transfected with a mutant MLP1 construct (three serine residues in the effector domain replaced with aspartate residues) showed a significant decrease in EnNaC activity compared to cells transfected with wildtype MLP1. Compared to vehicle treatment, mAoEC treated with the PIP2 synthesis blocker wortmannin showed less colocalization of EnNaC and MLP1. In other experiments, Western blot and densitometric analysis showed a significant decrease in MLP1 and caveloin-1 protein expression in mAoEC treated with wortmannin compared to vehicle. Finally, wortmannin treatment decreased sphingomyelin content and increased membrane fluidity in mAoEC. Taken together, our results suggest constitutive phosphorylation of MLP1 attenuates the function of EnNaC in aortic endothelial cells by a mechanism involving a decrease in association with MLP1 and EnNaC at the membrane, while deletion of PIP2 decreases MARCKS expression and overall membrane fluidity.

## Introduction

The amiloride sensitive epithelial sodium channel (ENaC) plays a rate-limiting role in the reabsorption of sodium in the distal segment of the nephron, colon, and lung airways (1). Amiloride sensitive sodium channels are also expressed in the vascular endothelium and these endothelial sodium channels (EnNaC) play an important role in regulating stiffness of the endothelium (2, 3). Like ENaC, membrane expression of EnNaC is stimulated by aldosterone (4) and EnNaC activity is inhibited by amiloride (4, 5). Conversely, there are some differences in the regulation of renal ENaC and endothelial EnNaC. Korte et al demonstrated a feedforward activation of EnNaC by sodium that contrasts the feedback inhibition of ENaC in the kidney (6).

Natriuretic peptides have been shown to inhibit ENaC activity in the kidney. A previous study showed activation of the clearance receptor for the natriuretic peptides (NPR-C) is coupled to the inhibition of EnNaC activity in human aortic endothelial cells (7). Downregulation of NPR-C gene expression and protein function has been observed in multiple disease models. For example, the deletion of the NPRC gene (*NPR3)* was shown to result in an increase in the severity of acute lung injury in obese mice (8). NPR3 down regulation was also demonstrated in a rabbit model of aortic coarctation (CoA) and in upstream aortic tissue from CoA patients (9).

Previous studies suggested bioactive lipids and myristoylated alanine-rich c-kinase substrate (MARCKS) enriched in extracellular vesicles (EVs) increase EnNaC activity in human aortic endothelial cells (10). Accumulating evidence suggests lipids play a role in regulating stiffness. An accumulation of ceramide has been shown to induce inflammation and mitochondrial oxidative stress (11). Another study showed oxidatively modified low density lipoprotein induces endothelium stiffening by causing a redistribution of lipid rafts (12).

The goals of this study were to identify specific amino acid residues within the effector domain of MARCKS protein that are involved in the regulation of EnNaC activity and determine whether depletion of lipid rafts affects membrane fluidity and the association between NPRC, EnNaC, and MARCKS protein in aortic endothelial cells.

## Materials and Methods

### Cell culture and treatments

Mouse aortic endothelial cells (mAoEC) were purchased from Cell Biologics (Cat No. is C57- 6052). The cells were maintained in endothelial cell medium (Cat No. M1168) and media exchanges were performed every 3 days. Only cells with a passage number of less than 9 was used for experiments. Cells were treated with wortmannin (Sigma-Aldrich, St. Louis, MO), USA) or DMSO (Sigma).

### Transient transfections of mutant and wild-type MARCKS constructs

mAoEC were transfected at 70% confluency with three mutant MARCKS/MLP1 constructs or a wild-type MARCKS/MLP1 construct (13) using the Xfect Transfection Reagent while following the manufactures instructions.

### Bicinchoninic colorimetric assay

Cells were scraped in ice cold mammalian protein extraction reagent (MPER) (ThermoFisher Scientific, Waltham, MA, USA). Cell lysates were sonicated on ice twice for 3 second intervals before being diluted 1:10 in type 1 water. Eight BCA standards were prepared from serial dilutions of a 2mg/ml stock solution of BSA (Sigma Aldrich) and each cell lysate sample was added to separate wells of a 96 well plate. A mixture of BCA reagents A and B (ThermoFisher Scientific) were added to each standard or sample well. After a 5-minute incubation the plate was read for absorbance at 570nm (Tecan plate reader) equipped with Magellan software. The concentrations of each sample were determined by linear regression.

### SDS-PAGE, Western blotting, and Densitometric Analysis

Each sample was prepared using 0.2 µm filtered ultrapure 1X PBS (Corning) and 50 µg total protein was loaded onto separate lanes of 4-20% tris glycine gels (ThermoFisher Scientific). The samples were resolved by electrophoresis using a Criterion apparatus (BioRad; Hercules, CA, USA). Next, the proteins were transferred to nitrocellulose membranes (ThermoFisher Scientific) using a Criterion apparatus (BioRad) and then stained with Ponceau stain for subsequent assessment of lane loading. The membranes were blocked with 5% non-fat milk prepared in 1X TBS (BioRad) for 1 hour and then incubated with primary antibodies (Table 1) each prepared at a 1:1000 dilution in 5% BSA 1XTBS overnight at 4°C. Afterwards, the blots were washed and incubated with a 1:3000 dilution of goat anti-rabbit (BioRad) or anti-mouse secondary antibodies (Bio-Rad) prepared in blocking solution for 1 hour. After a series of washes with 1XTBS the blots were incubated with ECL (BioRad) for 7 minutes. The blots were developed on a BioRad imaging system. Densitometric analysis was performed using ImageJ software (IJ 1.53k).

**Table 1.** Antibodies used in this study.

| Antibody | Application | Source |
| --- | --- | --- |
| EnNaC alpha 59 | Western blotting/Immunofluorescence | Alli et al |
| Caveolin-1 | Western blotting | Cell Signaling Technology |
| NPR-C JAH84 | Western blotting | Alli and Gower |
| MARCKS | Western blotting | Abcam |
| MARCKS | Immunofluorescence | Santa Cruz |

### Single-channel patch clamp studies

Micropipettes were pulled using a two-stage puller (Narishige, Tokyo, Japan) from filamented borosilicate glass capillaries (World Precision Instruments, Sarasota, FL USA). mAoEC were cultured on glass coverslips and transfected with wild-type or mutant MARCKS constructs. The cells were patched for EnNaC activity using the cell attached configuration 48 hours after transfection as previously described by our group (14). After applying different voltages to the cells an I-V curve was plotted and the conductance of the channel was calculated.

### Immunofluorescence microscopy

Mouse aortic endothelial cells were cultured on 35 mm glass bottom dishes (Mattek; Ashland, MA, USA) and incubated wortmannin or vehicle. The cells were washed with 1XPBS and then fixed for 10 minutes with ice cold methanol/acetone (1:1 v/v). After two washes with 1X PBS, the cells were incubated in 2.5% horse serum for 20 minutes. Next, they were incubated with primary antibodies against MARCKS and EnNaC alpha (Table 1) prepared in ultrapure 1X PBS at a 1:200 dilution for 45 minutes. The cells were then washed twice with 1X PBS and incubated with secondary antibodies (AF488 Cat No. A21206 and AF568 Cat No. A10037, Invitrogen).

Finally, after two more washes with 1X PBS, the cells were cover-slipped with antifade mounting media (Abcam) along with DAPI (Vector Laboratories Inc) and imaged using an Olympus microscope.

### Sphingomyelin Assay

Sphingomyelin levels in mAoECs after treatment with vehicle or wortmannin was determined by a sphingomyelin assay kit (abcam, ab138877) while following manufactures instructions.

### Membrane Fluidity assay

A membrane fluidity assay (abcam, ab189819) was performed while following the manufacturer’s instructions. Briefly, 5,000 mAoECs were plated per well of a clear bottom 96- well plate-coated. Next, the cells were labeled at 25°C for 30 minutes while being protected from light. Three washes with perfusion buffer were performed to remove any unincorporated pyrenedecanoic acid. The ratio of fluorescence at 460 nm and 405 nm was plotted after the background fluorescence was subtracted from each sample.

### Statistical Analysis

SigmaPlot software version 15 (Jandel Scientific, San Rafael, CA, USA) was used to perform a Student t-test to make comparisons between two groups or a one way ANOVA to compare more than two groups. A p-value <0.05 was considered statistically significant.

## Results

### Transient overexpression of a MARCKS mutant decreases EnNaC activity in mAoEC

The subcellular localization and function of MARCKS/MLP1 protein is dependent on amino terminus myristoylation, phosphorylation of serine residues within the basic effector domain of the protein, and electrostatic interactions between this domain and the inner leaflet of the plasma membrane. To investigate for the first time whether the activity of EnNaC in mAoECs is regulated by serine residues within the effector domain of MARCKS/MLP1 and/or myristoylation of the protein we transiently overexpressed mAoECs with various constructs including an S3A construct consisting of three serine residues in the effector domain being replaced with alanine residues, an S3D construct consisting of three serine residues in the effector domain being replaced with aspartate residues, or a GA construct consisting of the glycine residue in the myristoylation domain being replaced by an alanine residue. Next, mAoEC overexpressing each construct were patched and changes in single-channel characteristics of endogenous EnNaC were analyzed. As shown in Figure 1, mAoEC overexpressing the S3D construct compared to the wild-type construct resulted in a decrease in EnNaC activity mainly at the level of N, number of channels in a patch. Conversely, overexpression of the GA construct compared to the wild-type construct in mAoEC resulted in an increase in the number of channels in a patch and overall EnNaC activity (Figure 1). EnNaC activity and the number of channels in a patch were comparable in cells mAoEC overexpressing the S3A construct and mAoEC overexpressing the wild-type construct (Figure 1).

**Figure 1.**
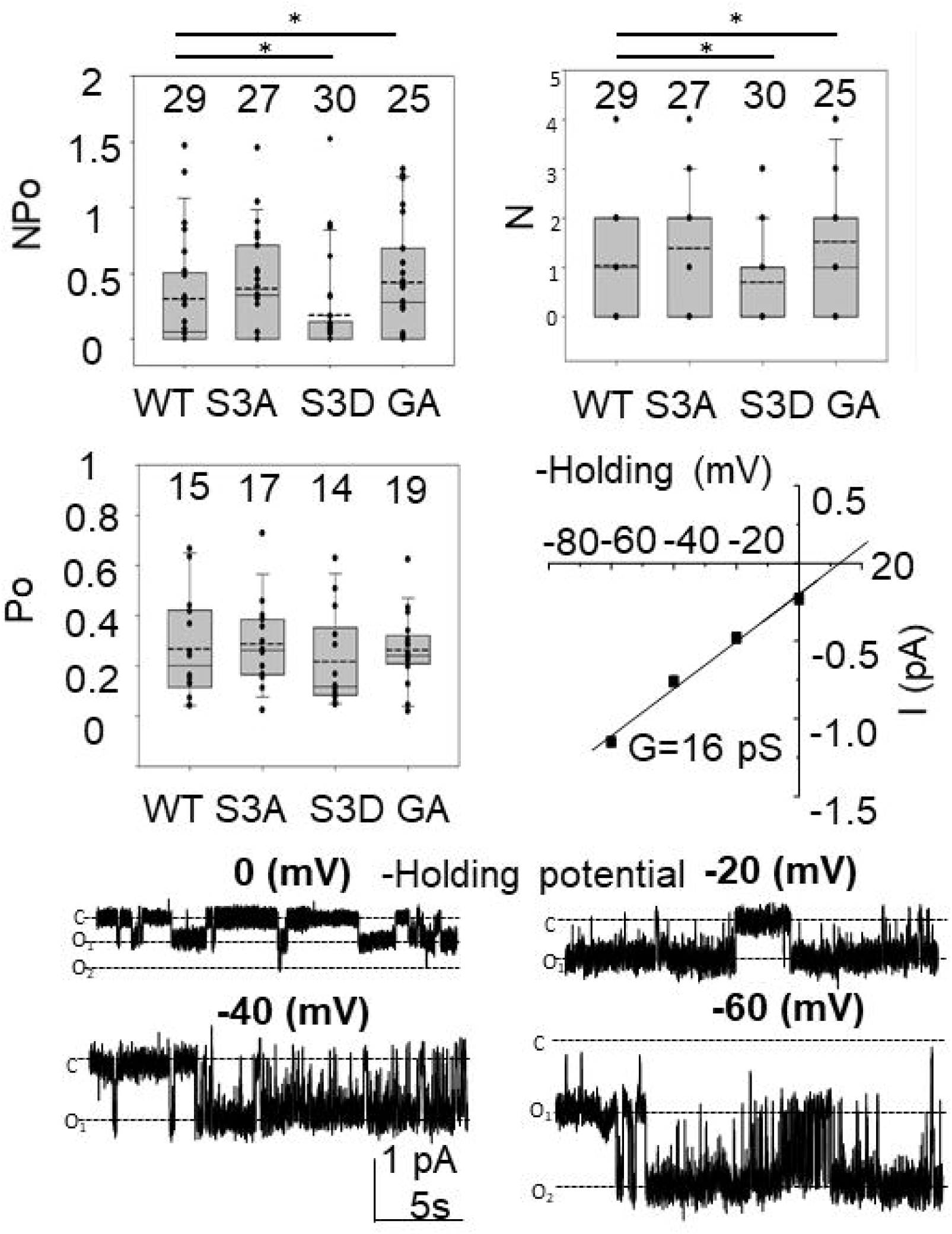
EnNaC activity in mAoEC transiently transfected with mutant or wild-type MARCKS constructions. After transient transfection of mAoEC the cells were patched for EnNaC activity by the single-channel patch clamp technique. A. Summary plot of EnNaC activity, NPo, B. Summary plot of the number of channels in a patch, N, C. Summary plot of the open probability of the channel, Po, D. Current voltage plot shown the conductance of EnNaC in mAoEC, E. Representative single channel recordings of EnNaC in mAoEC. The numbers at the top of the individual plots represent the number of patches per group. * represents a p-value <0.05.

### Wortmannin treatment attenuates MARCKS and EnNaC expression in mAoEC

Protein expression of both MARCKS and ENaC at the membrane is dependent on their interaction with the anionic phospholipid phosphate phosphatidylinositol 4,5-bisphosphate (PIP2). In renal epithelial cells of the distal nephron and collecting duct MARCKS sequesters PIP2 and increases its concentration in close proximity to ENaC to maintain the channel in an open confirmation. Here, we investigated for the first time whether depletion of PIP2 by wortmannin treatment affects MARCKS and EnNaC protein expression and their colocalization in mAoEC. Immunofluorescence microscopy studies showed less MARCKS and less EnNaC expression in mAoEC treated with wortmannin compared to vehicle (Figure 2). In addition, there is less colocalization of MARCKS and EnNaC in mAoEC treated with wortmannin (10µM) compared to cells treated with vehicle (Figure 2).

**Figure 2.**
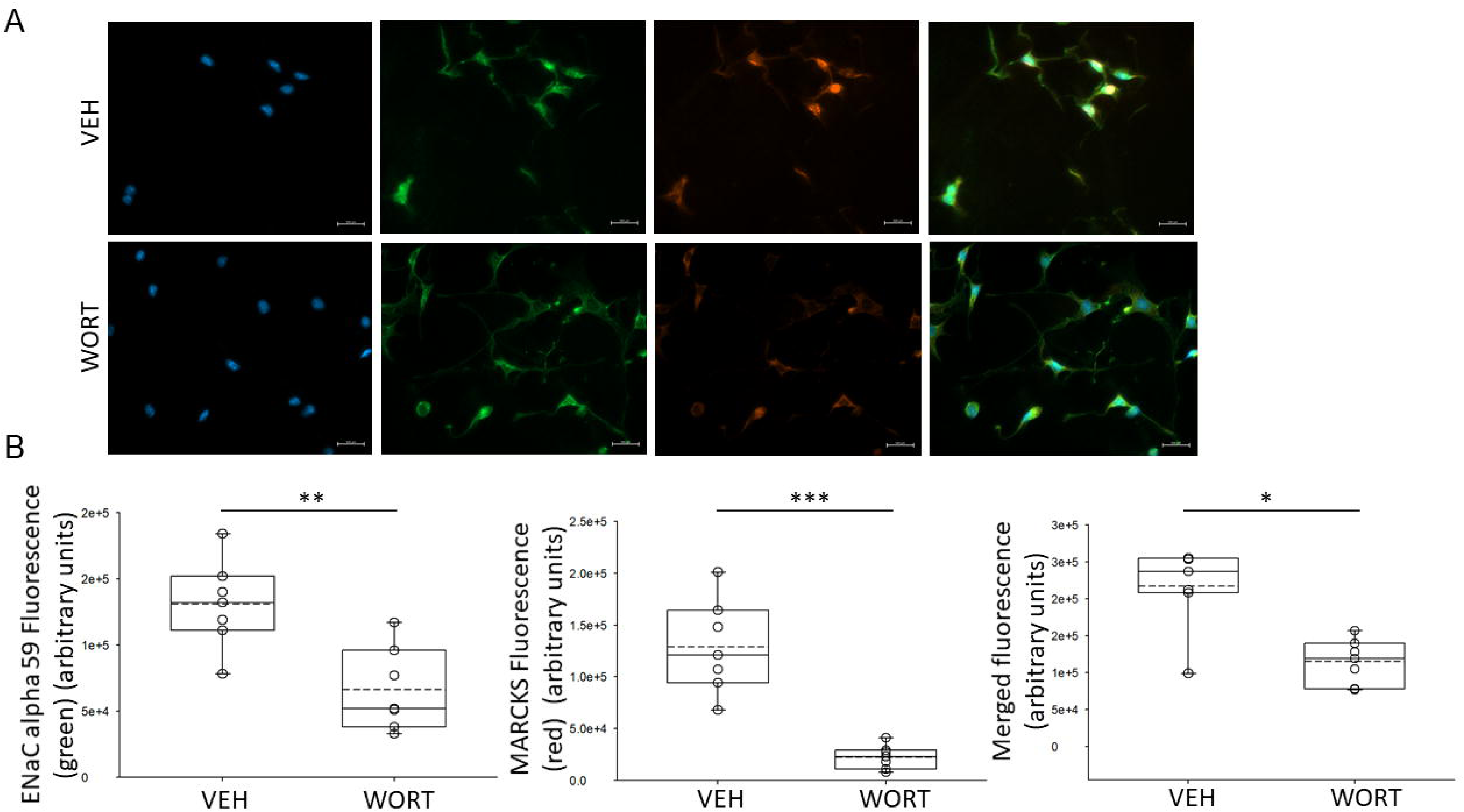
Immunofluorescence microscopy imaging and analysis of mAoEC cells treated with wortmannin or vehicle. A. Representative images showing MARCKS and EnNaC expression in mAoEC treated with wortmannin (WORT) or vehicle (VEH) for 8 hours. B. ImageJ analysis of the fluorescence intensity. Seven cells in each field were used for the analysis. **p-value<0.01, ***p-value<0.001.

### Wortmannin attenuates MARCKS and caveolin-1 density in lipid rafts fractions of mAoEC

There are several proteins that are associated with and enriched in lipid rafts. MARCKS and caveolin-1 proteins have been shown to be enriched in lipid rafts fractions. Here we investigated for the first time how wortmannin treatment (10µM) compared to vehicle treatment affects the expression of these proteins in membrane fractions of mAoEC. Wortmannin treatment significantly decreased MARCKS and caveolin-1 protein expression in membrane fractions of mAoEC compared to cells treated with vehicle (Figure 3).

**Figure 3.**
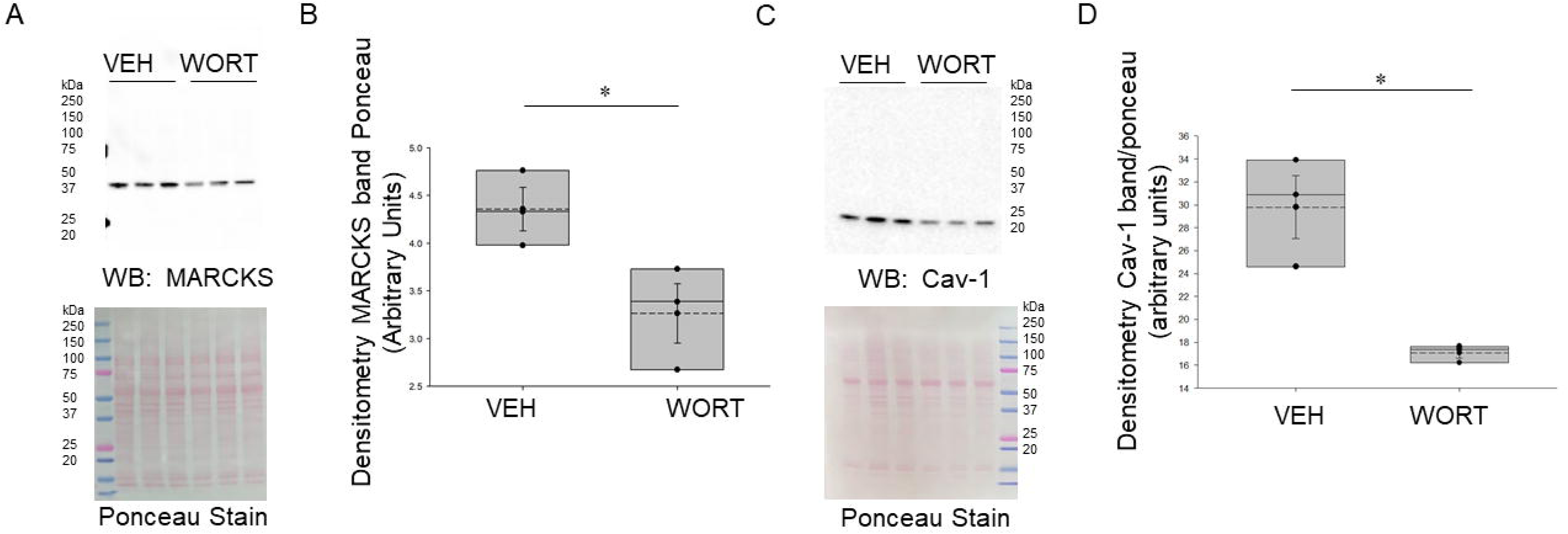
Western blot analysis of the expression of the lipid raft proteins MARCKS and Cav-1 in mAoEC treated with wortmannin or vehicle. A. Western blot of MARCKS in mAoEC treated with VEH or WORT. Ponceau stain used to assess lane loading. B. Densitometric analysis of the immunoreactive MARCKS band normalized to the Ponceau stain, C. Western blot of Cav-1 in mAoEC treated with VEH or WORT. Ponceau stain used to assess lane loading. D. Densitometric analysis of the immunoreactive Cav-1 band normalized to the Ponceau stain * represents a p-value<0.05. MARCKS, myristoylated alanine-rich C-kinase substrate; Cav-1, caveolin-1, mAoEC, mouse aortic endothelial cells; VEH, vehicle; WORT, wortmannin.

### Wortmannin reduces sphingomyelin concentration and increases membrane fluidity in mAoEC

Sphingomyelin has been reported to form cholesterol-dependent lipid raft microdomains at the plasma membrane. Therefore, we investigated whether wortmannin treatment (10µM) compared to vehicle treatment would affect sphingomyelin concentration in membrane fractions of mAoEC. As shown in Figure 4A, wortmannin treatment significantly decreased sphingomyelin concentration in mAoECs after treatment for 90 minutes. The amount of sphingomyelin in untreated mAoEC and vehicle treated mAoEC was comparable (Figure 4A). Since the density of lipid rafts are known to regulate membrane fluidity, we measured the effect of wortmannin treatment on membrane fluidity in mAoEC. Compared to vehicle treatment, wortmannin treatment increased membrane fluidity in mAoECs (Figure 4B).

**Figure 4.**
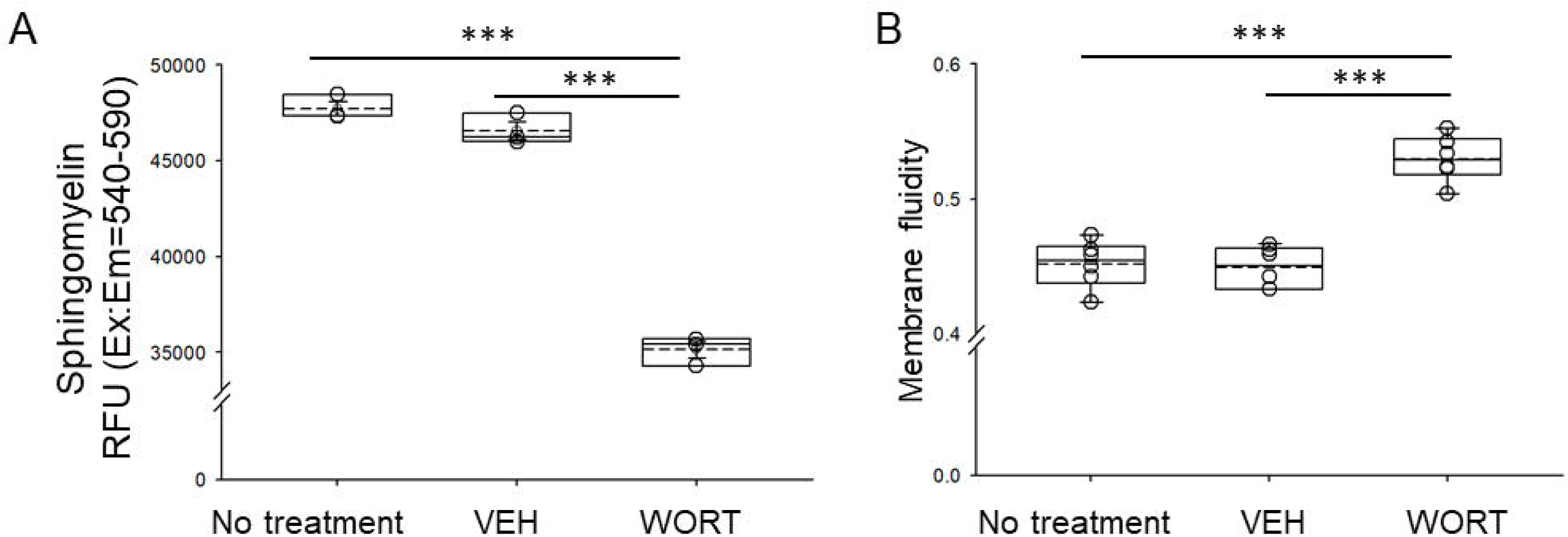
Assessment of membrane fluidity and sphingomyelin content in mAoEC treated with wortmannin or vehicle. A. Membrane fluidity in mAoEC treated with VEH or wort. B. Relative sphingomyelin content in mAoEC treated with VEH or wort. *** represents a p- value<0.001. mAoEC, mouse aortic endothelial cells; VEH, vehicle; WORT, wortmannin.

## Discussion

In the present study, mAoEC were used to investigate the role of specific amino acid residues within the effector domain and the myristoylation site of MARCKS/MLP1 in the regulation of EnNaC activity. Here we show mAoEC express a highly selective sodium channel (i.e. EnNaC) with a conductance of 16pS. In a previous study we reported the expression of a 25pS channel in human aortic endothelial cells (7). As expected, single channel characteristics of various ion channels could be different among species. For example, in *Xenopus* 2f3 cells the activity of the epithelial sodium channel (ENaC) is 5-7pS (15), but in mouse mpkCCD cells it is 17-20pS (14). Our electrophysiology data showed that the three serine residues within the effector domain of MLP1 plays an essential role in regulating EnNaC activity at the membrane of mAoEC. The S3D construct mimics constitute phosphorylation of MLP1 and presumably causes a loss of association between MLP1 at the plasma membrane since phosphorylation at these residues is thought to cause translocation of the protein to the cytoplasm. Compared to overexpression of full-length wild type MLP1, the overexpression of the S3D construct significantly decreased the number of channels in a patch and caused an overall decrease in EnNaC activity in these cells. On the other hand, overexpressing the GA construct in mAoEC surprisingly resulted in an increase in the number of channels in a patch and increase in EnNaC activity in mAoECs when compared to overexpression of the wild-type construct.

Overexpression of the GA construct would block myristoylation at the amino terminus of the protein and reduce hydrophobic interactions between MLP1 and the membrane.

Our group previously showed the function of EnNaC is negatively regulated by activation of the NPRC and inhibition of NPRC results in an increase in EnNaC activity (7).

Downregulation of NPRC protein expression has been shown to contribute to vascular modeling during spontaneous hypertension (21). Li et al showed overexpressing NPRC in vascular smooth muscle cells from spontaneously hypertensive rats resulted in a decrease in hypertrophy and hyperproliferation, and associated signaling pathways (21).

Increased activity of EnNaC results in adverse outcomes including, increased stiffness of the cortical actin cytoskeleton, reduced release of nitric oxide, enhanced inflammatory environment, increased oxidative stress, and augmented vascular permeability (16). An increase in EnNaC activity can be stimulated by a myriad of mechanisms including responses to organization of the actin cytoskeleton, hormones, extracellular vesicles, and lipids.

Some studies reported renal ENaC activity is positively regulated by the actin cytoskeleton (17–20). However, other studies reported the activity of renal ENaC is negatively regulated by actin cytoskeleton proteins (21, 22). In endothelial cells, an increase in EnNaC activity contributes to the stiffening of the cortical actin cytoskeleton (16). Transcriptional regulation of EnNaC by mineralocorticoid receptor mediated signaling in vascular endothelial cells was reported to dependent on the integrity of the cytoskeleton (23).

An increase in membrane expression and activity of EnNaC can contribute to increased endothelial stiffness. Jia et al showed osmotic minipump infusion of aldosterone in male and female mice induced endothelial stiffness and amiloride treatment reduced inward sodium currents in isolated mouse endothelial cells (2). A combination of mineralocorticoid receptor activation and a high salt diet was reported to activate EnNaC and induce arterial stiffening (23). Results from a different study indicated endothelial function is improved and aortic stiffness is reduced by a low dose of amiloride in female mice fed a high fat Western diet (24).

A previous lipidomic study by our group showed the conditioned media of human aortic endothelial cells contain extracellular vesicles enriched in sphingomyelins, phosphatidylcholine (PC), and lysophosphatidylcholine (LPC), phosphatidylethanolamine (PE), phosphatidylserine (PS) (10). A comparative study by our group showed mpkCCD cells produce two types of EVs in which the apical plasma membrane EVs are enriched in sphingomyelins while the basolateral plasma membrane EVs are enriched in PC, PG, PS, CL, and Cer (25). Phospholipids including sphingomyelins have been shown to form cholesterol-dependent raft microdomains (26, 27).

Our data of wortmannin decreasing sphingomyelin content in mAoEC and increasing membrane fluidity in these cells is consistent with the literature. Horváth et al showed methyl-β- cyclodextrin (MBCD) increases membrane fluidity (28). MBCD has been shown to disrupt lipid rafts by depleting cholesterol. Sphingomyelin, like cholesterol, is known to decrease membrane fluidity. Zager et al showed the addition of sphingomyelin to cultured human proximal tubule cells alleviated oxidant injury presumably by sphingomyelins effects on decreasing membrane fluidity (29). PIP2 is enriched in lipid rafts and is known to positively regulate MARCKS and ENaC function at the apical plasma membrane of distal tubule and collecting duct cells, and presumably prevents these proteins from degradation mechanisms. Here, we show depletion of PIP2 levels and organization of lipid rafts by wortmannin treatment decreased the association between MLP1 and EnNaC proteins in mAoEC while increasing membrane fluidity (Figure 5).

**Figure 5.**
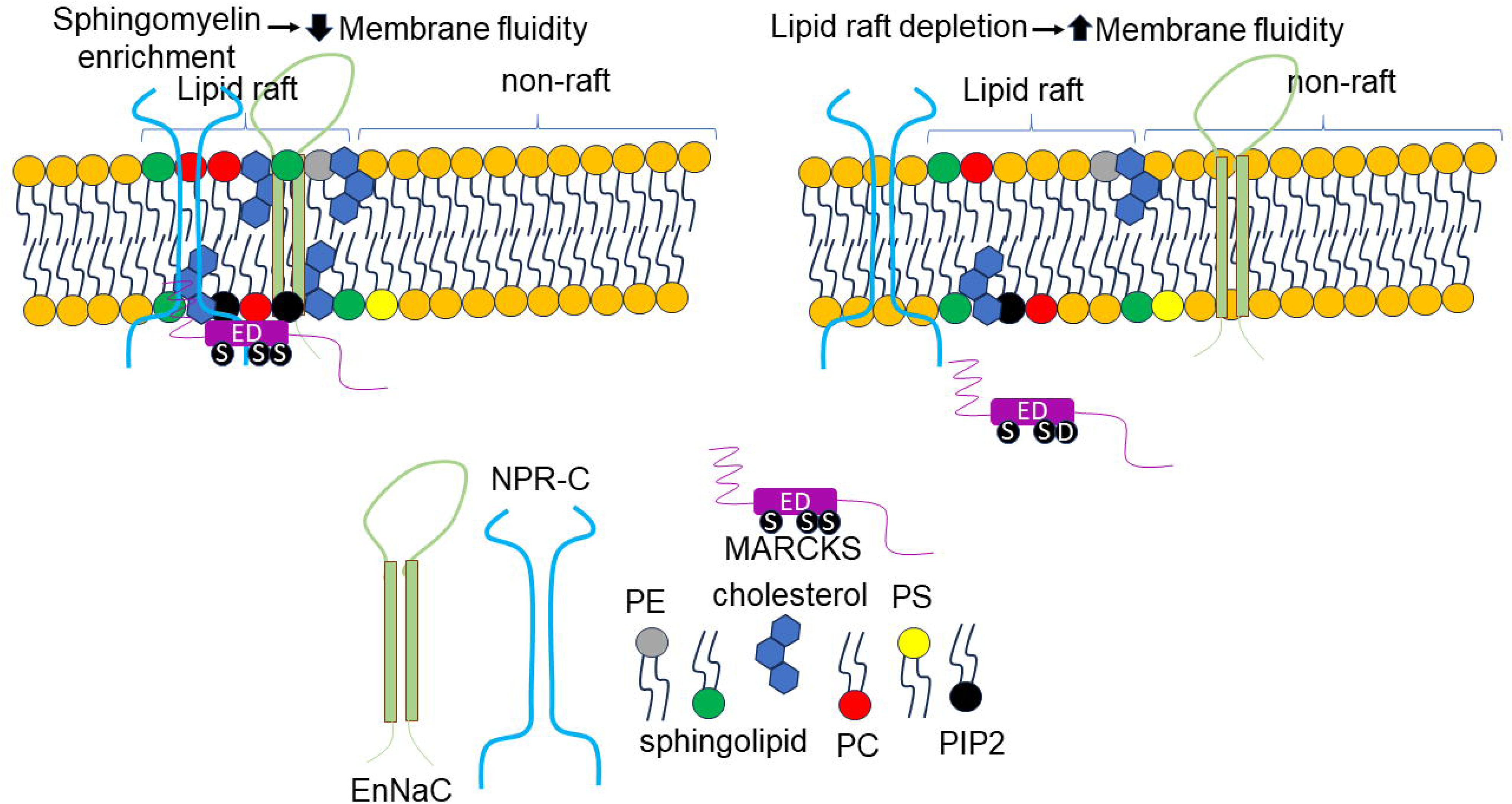
Proposed model of the role of the effector domain of MARCKS and lipid rafts in the crosstalk between NPRC and EnNaC. PE, phosphatidylethanolamine; PS, phosphatidylserine; PC, phosphatidylcholine; MARCKS, myristoylated alanine-rich c-kinase substrate

Although it is known NPRC activation can regulate EnNaC function in endothelial cells, we did not investigate whether phosphorylation of MLP1 affects the cross-talk between NPRC and EnNaC in endothelial cells. Another limitation of our study is that we did not investigate whether depletion of PIP2 and disruption of lipid rafts affects the interaction between NPRC and EnNaC. Additional experiments can be performed to address these knowledge gaps.

In summary, the data presented here are consistent with our hypothesis that MLP1 phosphorylation negatively regulates EnNaC activity and depletion of PIP2 decreases the association of MLP1 and EnNaC in mAoEC. Presumably, a decrease in PIP2 concentration in mAoEC increases membrane fluidity in these cells by disrupting the organization of lipids rafts.

## Acknowledgments

The authors thank Dr. Douglas Eaton for kindly providing us with the MLP1 mutant constructs and MLP1 wildtype construct used for the electrophysiology studies.

## Author Contributions

LY, NB, VLN, LK and AAA performed experiments and analyzed data. AAA provided supervision and wrote the manuscript. LY, NB, VLN, LK, JFL, Jr, and AAA edited and approved the final version of this manuscript.

